## Supplementary Notes, Table, Figures and Videos for "DeepCristae, a CNN for the restoration of mitochondria cristae in live microscopy images"

**Salomé Papereux[1,2,&], Ludovic Leconte[1,2,&], Cesar Augusto Valades-Cruz[1,2,&], Tianyan Liu[3], Julien Dumont[4], Zhixing Chen[3], Jean Salamero[1,2], Charles Kervrann[1,2], Anaïs Badoual[1,2,*]**

1 SERPICO Project Team, Centre Inria de l'Université de Rennes, F-35042 Rennes, France

2 SERPICO Project Team, UMR144 CNRS Institut Curie, PSL Research University, F-75005, Paris, France

3 College of Future Technology, Institute of Molecular Medicine, National Biomedical Imaging Center, Beijing Key Laboratory of Cardiometabolic Molecular Medicine, Peking-Tsinghua Center for Life Science, Academy for Advanced Interdisciplinary Studies, Peking University, Beijing 100871, China

4 CIRB Microscopy facility, Collège de France, UMR 7241 CNRS, Inserm U1050, Paris, 75005, France

[&] these authors contributed equally to the work

| **Supplementary Material** | **Title** |
|---|---|
| **Supplementary Note 1** | Materials and Methods. |
| **Supplementary Note 2** | Network Evaluation. |
| **Supplementary Table 1** | Dataset $D_{synt}$ and its division into training, validation and test sets. |
| **Supplementary Figure 1** | Illustrations of binary masks for areas of interest. |
| **Supplementary Figure 2** | Robustness of DeepCristae to mitochondria scale in LR images and limitations of use. |
| **Supplementary Figure 3** | Stability of image restoration by DeepCristae. |
| **Supplementary Figure 4** | Reliability of image restoration by DeepCristae. |
| **Supplementary Figure 5** | DeepCristae applied to AiryScan processed images and Lattice Light Sheet Microscopy. |
| **Supplementary Figure 6** | Cristae density evolution during fission in or out of lysosome contacts with mitochondria. |
| **Supplementary Figure 7** | Pipeline of the image patch sampling for the training step. |
| **Supplementary Figure 8** | Ablation study for different components of our method. |
| **Movie S1** | DeepCristae restoration enhances cristae resolution in live imaging using a Spinning-Disk confocal microscope equipped with Live-SR (related to Fig. 7e). |
| **Movie S2** | DeepCristae restoration enhances cristae resolution in live imaging using a Lattice Light Sheet microscope (related to Fig. 7f). |
| **Movie S3** | DeepCristae reveals 3D+time cristae morphology during endo/lysosome mitochondria interactions using a Spinning-Disk confocal microscope equipped with Live-SR (related to Fig. 8a). |
| **Movie S4** | DeepCristae reveals 3D+time cristae morphology during endo/lysosome mitochondria interactions using Lattice Light Sheet microscope (related to Supplementary Fig. 5d-e). |

### Supplementary material

#### Supplementary Note 1: Materials and Methods

#### 1.1 Cell culture and fluorescence labeling

The hTERT-immortalized RPE1 cells (Human Retinal Pigment Epithelial Cell) were purchased from ATCC (CRL-4000). 150 to 250 × 103 RPE1 cells were plated on glass bottom Petri dishes (µ-dish 35mm-High, 1.5H, Ibidi, GmbH, Gräfelfing, Germany) for STED, Spinning Disk (SD)-Live-SR or AiryScan confocals. For LLSM (Lattice Light Sheet Microscopy) imaging, cells were seeded 5 to 6 hours before image acquisition at 150 × 103 per well of a 6 well plate, containing 3 to 4 coverslips with a 5mm diameter, treated as previously described in Chen et al.[1]. Live cells were stained for mitochondria by diluting 4000× a 1mM stock solution of PKMITO-Orange[2] (Spirochrome AG, Switzerland and Genvivo Biotech, Nanjing, China) in DMEM/F12 medium for a final 62.5 nM, 15 minutes at 37° before image acquisition. For labeling of the endosomal pathway, cells were incubated with the CellMaskTM Plasma Membrane DeepRed probe at a 1/10,000 dilution of the stock solution prepared as indicated by the provider (Molecular Probes, Life Technologies, OR, USA) for 3 to 4 hours at 37°C allowing slow endocytosis and labeling the whole endo/lysosomal compartments, before incubation with PKMITO-OrangeTM, as above. To acquire images of lysosomes in living cells, the cells were incubated with SIR-lysosome (Spirochrome AG, Switzerland) for 2 hours at 37°C before imaging.

#### 1.2 Microscopies

**1.2.1 STED microscopy**. Image acquisitions were performed with a STEDYCON module (Abberior Instruments, Göttingen, Germany) mounted at the camera port of a Zeiss AxioObserver 7 Videomicroscope (Carl Zeiss Microscopy GmbH, Iena, Germany) with a Plan-ApoChromat DIC Oil ×100 objective (1.4 NA) or at the camera port of a TCS SP8 STED microscope (Leica, Manheim, Germany) with a HC PL APO C2S ×100 oil objective (1.4 NA) used in 2D mode.

For the creation of the 2D STED dataset $D_{synt}$ used to train DeepCristae, the 33 initial high-resolution (HR) STED images (denoted by $I_{HR}$ in Methods "Generation of the 2D STED dataset" and Fig. 1) were obtained as follows: depletion was performed with a 775 nm pulsed laser 1-7 ns at 80-100% (413 mW) at the objective lens position. Labeled mitochondria were imaged with excitation at a wavelength of 565 nm with nominal laser power adjusted at about 13% (3.15 µW) at the objective lens position. The time-gated fluorescence detection was done on a detector single photon counting avalanche photodiodes between 575-625 nm with a pinhole settled at 64 µm. Imaging was executed with 3 to 14 lines accumulation. Pixel dwell time was 10 µs and the pixel size of 25 × 25 nm.

For the pairs of real LR and HR 2D STED images (Results "Reliability of image restoration by DeepCristae"), other imaging parameters were used at high (HR, 100% laser depletion; 3-lines accumulation) and low (LR, depletion between 40 and 100%; 3 lines accumulation) resolutions. The manual transition from HR to LR mode takes about 30 s.

For 2D+time STED dataset, HR images were acquired mainly as above, with a systematic depletion at 100%, excitation adjusted at 7%, and 10-lines accumulation for emission, with two different pixel sizes (25 × 25 nm or 50 × 50 nm) when indicated. For LR 2D+time STED images (also referred as Fast STED images), depletion was reduced to 40-64%, and 3-line scans accumulated with a pixel size of 25 × 25 nm. The series are constituted of at least 10 timepoints each, with a duration of ~13 s/image and 3 to 6 s/image for HR and LR (Fast) STED, respectively.

**1.2.2 Live-SR microscopy**. Acquisition was performed using a spinning disk confocal based on a Ti2-Eclipse inverted microscope (Nikon, Tokyo, Japan) equipped with W1 confocal head (Yokogawa, Tokyo, Japan), an sCMOS camera (Prime95B,Teledyne Photometrics, Tucson, Arizona, USA), a stage top incubator (Tokai Hit, Hokkaido, Japan), an automated piezo stage (Nano z500, Mad City Lab, Madison, WI, USA) and interfaced with a super resolution Live-SR module (GATACA systems, Massy, France) based on optically demodulated structured illumination technique. Cells were scanned incrementally through ~4 µm Z-stacks in 200 nm step at 30 ms/frame, using a 100×, CFI Plan Apo NA 1.4 oil objective (Nikon, Tokyo, Japan). Pixel size is 65 × 65 nm. Excitations were achieved at 561 and 633 nm (150mW and 100mW) diode lasers (GATACA systems, Massy, France), at low illumination of the sample adjusting the acousto-optic tunable filter transmittance at ~20% in the Live-SR mode for 561 Ex/ 610 Em and at 14% max for 633 Ex/700 Em. The acquisition and reconstruction of the Live-SR module is performed with Metamorph 8.6 software (Molecular Devices, San Jose, California, USA). Image datasets are constituted of 3D+time series for a duration between 3 and 4 minutes (15-20 planes/stack; acquisition times/stack of 20 planes for 2 channels: ~4 s). Maximum intensity projections were generated using ImageJ/Fiji 1.53t. Before DeepCristae restoration, all 2D images were transformed using ImageJ/Fiji "Bio-formats" (https://github.com/ome/bio-formats-documentation/). Endosomal/lysosomal pathway labeling (CellMask™ Plasma Membrane DeepRed) corresponding datasets are first denoised using ND-SAFIR[3] software (https://gitlab.inria.fr/serpico/ndsafir_bin) before deconvolution using Richardson-Lucy (RL) algorithm. Richardson-Lucy deconvolution was performed using *deconvlucy* from MATLAB2019b imaging processing toolbox. Napari, a multi-dimensional image viewer for Python (https://github.com/napari/napari), was used for 3D rendering while for color 3D rendering, ImageJ/Fiji "temporal-color code" function was used. Related Supplementary Movies S1 and S3 were generated using napari and napari-animation plugin (https://github.com/napari/napari-animation) and the final film montages using DaVinciResolved 18.1.4 (https://www.blackmagicdesign.com/products).

**1.2.3 LLSM microscopy.** Image acquisition was performed on a commercialized version of a previously described setup from 3i (Denver, USA). Cells were scanned incrementally through a 20 µm long light sheet, in 600 nm steps using a fast piezoelectric flexure stage (equivalent to ≃ 325 nm voxel size in z axis after image realignment, respecting the 32.8° angle for the detection objective position). Data were imaged using sCMOS camera (Orca-Flash 4.0; Hamamatsu, Bridgewater, NJ). Excitation was achieved with 561 nm or 642 nm (MPB Communications, Montreal, Canada) diode lasers at ~5% acousto-optic tunable filter transmittance, with 50 mW nominal power. Excitation is done through a water-dipping 28.6× objective ($\mathrm{NA} = 0.7$, working distance 3.74 mm) and detection via a Nikon CFI Apo LWD 25× water-dipping objective ($\mathrm{NA} = 1.1$), completed with a 2.5× tube lens, to obtain a final pixel size of 104 × 104 nm. Lattice light-sheet imaging was performed using an excitation pattern of outer NA equal to 0.55 and inner NA equal to 0.493. Composite volumetric datasets were obtained using ≃ 5 ms exposure/optical planes/channel at a time resolution of 1 to 1.3 second per total cell volume of 60 planes. Double labeled images were acquired on two separated cameras, using a dichroic mirror with an edge at 660 nm. 3D+time series were acquired within 2 to 4 minutes. Acquired data were deskewed, a necessary step to realign the image frames, using LLSpy, a python library (copyright to T. Lambert, Harvard Medical School, Boston, USA; https://github.com/tlambert03/LLSpy) and deconvolved using *cudaDeconv* (copyright to Lin Shao et al, HHMI https://github.com/dmilkie/cudaDecon), included in LLSpy. Before DeepCristae restoration, all 2D images were transformed using ImageJ/Fiji "Bio-formats" (https://github.com/ome/bio-formats-documentation/). For Endosomal/lysosomal pathway labeling (CellMaskTM Plasma Membrane DeepRed), LLSM images were denoised using ND-SAFIR[3] software (https://gitlab.inria.fr/serpico/ndsafir_bin) before deskewing and deconvolution, as above. Napari, a multi-dimensional image viewer for Python (https://github.com/napari/napari), was used for 3D rendering while for color 3D rendering, ImageJ/Fiji "temporal-color code" function was used. Maximum intensity projections were generated using ImageJ/Fiji 1.53t. Related Supplementary Movies S2 and S4 were generated using napari and napari-animation plugin (https://github.com/napari/napari-animation) and the final film montages using DaVinci Resolved 18.1.4 (https://www.blackmagicdesign.com/products/davinciresolve/).

**1.2.4 LSM980 AiryScan 2.** Image acquisition was performed with a confocal laser scanning microscope ZEISS LSM 980 (Carl Zeiss AG, Oberkochen, Germany) equipped with an Airyscan 2 detection unit and 2 PMTs and GaAsp detector array allowing spectral imaging. Cells were scanned incrementally through ~4 µm Z-stacks in 250 nm step, using high NA Plan-Apochromat 63×/1.4 Oil objective. Detector gain and pixel dwell times were adjusted for each dataset keeping them at their lowest values in order to avoid saturation and bleaching effects. Excitations were achieved at 561 (10mW) with diode lasers (Zeiss system) and adjusting the acousto-optic tunable filter transmittance at 13% for 561 Ex/ 610 Em. Image datasets are constituted of 3D+time series for a duration between 1.5 to 2.5 minutes (15 planes/stack; acquisition times/stack for 1 channel, 17 to 30 s). The reconstruction of the AiryScan images was performed based on the values automatically determined by the Zen software (Carl Zeiss AG, Oberkochen, Germany). Before DeepCristae restoration, all 2D images were transformed using ImageJ/Fiji "Bio-formats" (https://github.com/ome/bio-formats-documentation/).

**1.3 Measurements**

Line profiles plots were extracted with ImageJ/Fiji1.53t and then fitted to a Gaussian model using our Fiji macros (doFWHM_distanceCristae_Per_Image_ROIS.ijm and doFWHM_distanceCristae_Per_IntensityProfile_ROIS.ijm), which are available on Figshare (DOI: 10.6084/m9.figshare.26940892). Note that these macros were derived from the Fiji macro FWHM.ijm (https://gist.github.com/lacan/45f865b5a38d7c3a96a7cd8b25923407#file-fwhm-ijm) created by Olivier Burri and Romain Guiet from the BioImaging & Optics Platform (BIOP). Full width at half maximum (FWHM) is estimated from the fitting results. Student t-test and Fisher-test were performed using GraphPad Prism 9. Evaluation metrics based on *NRMSE*, *PSNR* and *SSIM* were computed using our Jupyter notebook demo_metrics.ipynb available at https://github.com/SalomePx/DeepCristae/notebooks.

**Supplementary Note 2: Network Evaluation**

**2.1 Image quality assessment**

To evaluate the performance of our proposed method and compare it to state-of-the-art methods, we used the following metrics that are widely used in image restoration. Let us denote $y, \hat{y} \in R^{W \times L}$, the reference image and its corresponding prediction, respectively. They are defined over the grid $\Omega$ of size $W \times L$.

**NRMSE.** The normalized root mean square error between a reference image $y$ and a prediction $\hat{y}$ is defined by

$$NRMSE(y, \hat{y}) = \frac{\sqrt{MSE(y, \hat{y})}}{y_{max} - y_{min}}, \tag{5}$$

where $y_{max}$ and $y_{min}$ are the maximum and minimum values of $y$, respectively, and

$$MSE(y, \hat{y}) = \frac{1}{W \times L} \sum_{(i,j) \in \Omega} \left(y(i,j) - \hat{y}(i,j)\right)^2. \tag{6}$$

A value of zero of the *NRMSE* indicates a perfect restoration.

Since the restored image $\hat{y}$ (predicted by DeepCristae or any comparative method) and the corresponding ground truth image y have different dynamic range of intensity value, we first normalize the images to a common range. To that end, we follow the approach adopted by CARE[4], namely is solved the 2D ordinary least squares (OLS) regression between the normalized ground truth $y_{norm}$, using the percentile-normalizer described in Eq. (2) (Methods), and the transformed prediction $\alpha\hat{y} + \beta$ . More precisely, we solve $\alpha_{OLS}, \beta_{OLS} = argmin_{\alpha,\beta}\ MSE\ (y_{norm}, \alpha\hat{y} + \beta)$ . Finally, all *NRMSE* metrics were computed using this mapping of intensity value, that is to say as $NRMSE(y_{norm}, \widehat{y_{OLS}})$ with $\widehat{y_{OLS}} = \alpha_{OLS}\hat{y} + \beta_{OLS}$, allowing to obtain more relevant metrics scores without modifying the semantic meaning of the prediction.

**PSNR.** The peak signal-to-noise ratio expressed in dB is given by

$$PSNR(y, \hat{y}) = 10 \cdot log_{10}\left(\frac{MAX_y}{MSE(y, \hat{y})}\right) \tag{7}$$

where *MSE* is defined by Eq. (6) and $MAX_y$ is the maximum possible pixel value of $y$. In our case, $MAX_y = 255$ as we normalize $y$ and $\hat{y}$ between 0 and 255 before calculating the *PSNR*. The higher the *PSNR* value, the better the restored image quality.

**SSIM.** While the NRMSE and PSNR compare images pixelwise, the structural similarity index[5] considers image degradation as perceived changes in structural information variation. It takes into account the luminance *l*, the contrast $c$ and the structure $s$ of the image through the following formula

$$SSIM(y, \hat{y}) = l(y, \hat{y}) \cdot c(y, \hat{y}) \cdot s(y, \hat{y}). \tag{8}$$

The *SSIM* index ranges between -1 and 1. A value of 1 means that the two compared images are very similar, whereas a value of -1 indicates a significant visual difference. Before calculating the SSIM, we first normalize $y$ and $\hat{y}$ between 0 and 255.

**Customized metrics.** In the context of mitochondria cristae restoration, these three metrics are not sufficient to give a relevant measure of the quality of the cristae restorations, but only of the image as a whole. Indeed, the studied images contain only few pixels of cristae and thus have too little weight in these metrics compared to the background. To overcome this issue, we forced these metrics to focus their evaluation on the mitochondria pixels. We call these mitochondrial metrics $NRMSE_{mito}$, $PSNR_{mito}$, and $SSIM_{mito}$. Their formula is still given by Eqs. (5), (7) and (8), respectively, but instead of being computed on the whole image, they are computed over the set $\Omega_{mito}$ which contains the mitochondria pixels. Note that in the code, $\Omega_{mito}$ can be obtained either automatically by thresholding the Z-score map (Methods, Eq. (4)) or from a binary mask provided by the user. All the results presented in this paper were obtained from provided binary masks obtained semi-automatically by thresholding the Z-score maps plus manual refinements (Supplementary Fig. 1, second column). To go even further and accurately assess cristae restoration, we similarly introduced the so-called cristae metrics $NRMSE_{cristae}$, $PSNR_{cristae}$ and $SSIM_{cristae}$. They are computed over the set $\Omega_{cristae}$ which contains the pixels of the mitochondria cristae. For our experiments, $\Omega_{cristae}$ was obtained by manually annotating some cristae, using ImageJ/Fiji, in all the HR STED test images of $D_{synt}$ (Supplementary Fig. 1, third column).

### 2.2 Implementation details

**2.2.1 DeepCristae**. For all experiments mentioned in the main text as well as in the Supplementary material, we trained DeepCristae once (1,365,473 trainable parameters), unless specified, using the training set of $D_{synt}$ on a single NVIDIA A100 40 Go PCIe for 25 epochs with a batch size 8. These 25 epochs result from an early stopping based on the evolution of the loss on the validation set, with a patience of 15. We used the Adam optimizer with an initial learning rate of $4 \cdot 10^{-4}$ and decayed by 0.5 every 10 epochs if the model does not improve. DeepCristae was implemented in Python (TensorFlow version 2.11) and is freely available as an open-source software in our GitHub site (https://github.com/SalomePx/DeepCristae) as well as in the BioImageIT platform[6].

**2.2.2 Comparative methods.**

**Conventional methods (Richardson-Lucy[7,8], Wiener[9], SPITFIR(e)[10]).** We used the implementation provided by BioImageIT[6] with the default parameters, except for the padding that we set to True.

**CARE**[4]**.** We used the implementation available at https://github.com/CSBDeep/CSBDeep. We used the same parameters as the original model set by the authors, with the *MAE* loss to minimize (332,641 trainable parameters).

**ESRGAN**[11]**.** We used the code available at https://github.com/wonbeomjang/ESRGAN-pytorch. Unless otherwise specified, we used the same parameters as the original model set by the authors. We set the number of convolutional layers for feature extraction to 32; the scaling factor is set to 1 so that the input and output of the network have the same size. For the early stopping, we used a patience of 30. It finally has a total of 3,340,705 trainable parameters.

**RCAN**[112,13]**.** We took the original RCAN architecture, using their code (available at https://github.com/yulunzhang/RCAN), and reimplemented it with Tensorflow version 2.11. We used the same parameter described in the paper[13]. We set the number of Residual Groups (RG) to G = 5 in the Residual in Residual (RIR) structure; in each RG, the RCAB number is set to 3; the number of Convolutional Layers (CL) in the shallow feature extraction and RIR structure is C = 32; the CL in channel-downscaling has C/r = 4 filters, where the reduction ratio r is set to 8; the upscaling module at the end of the network is omitted because network input and output have the same size in our case. It finally has 1,339,509 parameters. The *MAE* is used as the training loss. In the original RCAN paper, a small patch with size 48×48 pixels is used for training. By contrast, we used a larger patch size (128×128 pixels), also convenient with the $D_{synt}$ dataset. We used the same stop condition and learning rate as DeepCristae.

**SRResNet**[14]**.** We took the original SRResNet architecture (displayed at https://github.com/twtygqyy/pytorch-SRResNet) and re-coded it with Tensorflow version 2.11. It is a succession of 16 residual blocks without upsampling, because network input and output have the same size in our case. The loss used for training is the *MSE*, the same as in the paper. It has a total of 1,233,281 trainable parameters.

#### 2.3 Ablation study

In order to highlight the individual contribution of key components of our method, we performed an ablation study. We conducted the following experiments:

- Depth 2: reduction of the number of down and up sampling to 2 in our network;
- $SCoP_{1,1}$: substitution of $\gamma = (1,4)$ by $\gamma = (1,1)$ in our SCoP loss (Methods, Eq. (1)), which is equivalent to the structural dissimilarity (*DSSIM*) loss;
- $SCoP_{1,6}$: substitution of $\gamma = (1,4)$ by $\gamma = (1,6)$ in our SCoP loss (Methods, Eq. (1));
- *MAE*: substitution of our *SCoP* loss by the *MAE* loss;
- No DA: removal of the data augmentation in the creation of our patches;
- Grid: substitution of our patch sampling method (Methods) by a simple grid patch sampling.

Each change in our method is performed separately. The table in Supplementary Fig. 8a, based on the *NRMSE*, *PSNR* and *SSIM* metrics, shows that our depth of 3 plays a major role in the performance of our network. With regard to SCoP losses, we also observe that when the second component of $\gamma$ (weighting the contribution of the background in the loss) is too large, such as $SCoP_{1,6}$, performance begins to decline; some granularity appears in the mitochondria. Otherwise, all other components of our method seem to play an important role in an equal measure. However, no significant conclusions can easily be drawn from this table. Hence, we also measured cristae widths for 155 cristae from the test set of $D_{synt}$ by fitting line profiles (Supplementary Fig. 8b) to a Gaussian model and measuring the FWHM. After restoration by the different methods of the ablation study, the best results in terms of mean, standard deviation and detection are obtained with our model (Supplementary Fig. 8c). This improvement is statistically relevant as confirmed by the results shown in Supplementary Fig. 8d.

### Supplementary Tables

**Supplementary Table 1.** Dataset $D_{synt}$ and its division into training, validation and test sets.

| | Number of images |
|---|---|
| Original training set | 24 |
| Data augmentation (x4) | 96 |
| Patch sampling (~ 19/image) | 1824 |
| Training set (80%) | 1459 |
| Validation set (20%) | 365 |
| Test set | 9 images sampled into 26 ROIs |

### Supplementary Figures

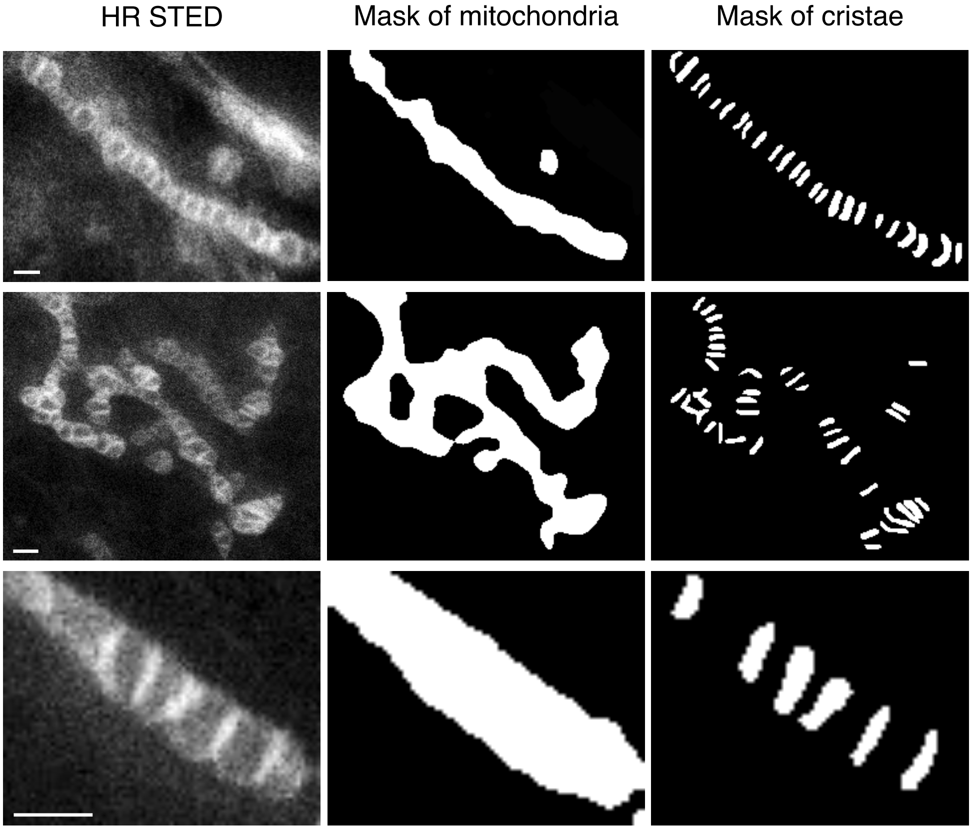


**Supplementary Figure 1. Illustrations of binary masks for areas of interest.** *First column*: three high-resolution (HR) STED images of $D_{synt}$. *Second column***:** corresponding binary masks of mitochondria obtained by thresholding the Z-score maps (see Methods, Eq. (4)) plus manual refinements. For each image, the white pixels define the set $\Omega_{mito}$ on which are computing the metrics $NRMSE_{mito}$, $PSNR_{mito}$ and $SSIM_{mito}$. *Third column*: corresponding binary masks of cristae obtained from manual annotations. For each image, the white pixels define the set $\Omega_{cristae}$ on which are computing the metrics $NRMSE_{cristae}$, $PSNR_{cristae}$ and $SSIM_{cristae}$. Pixel size: 25 nm. Scale bar: 0.5 µm.

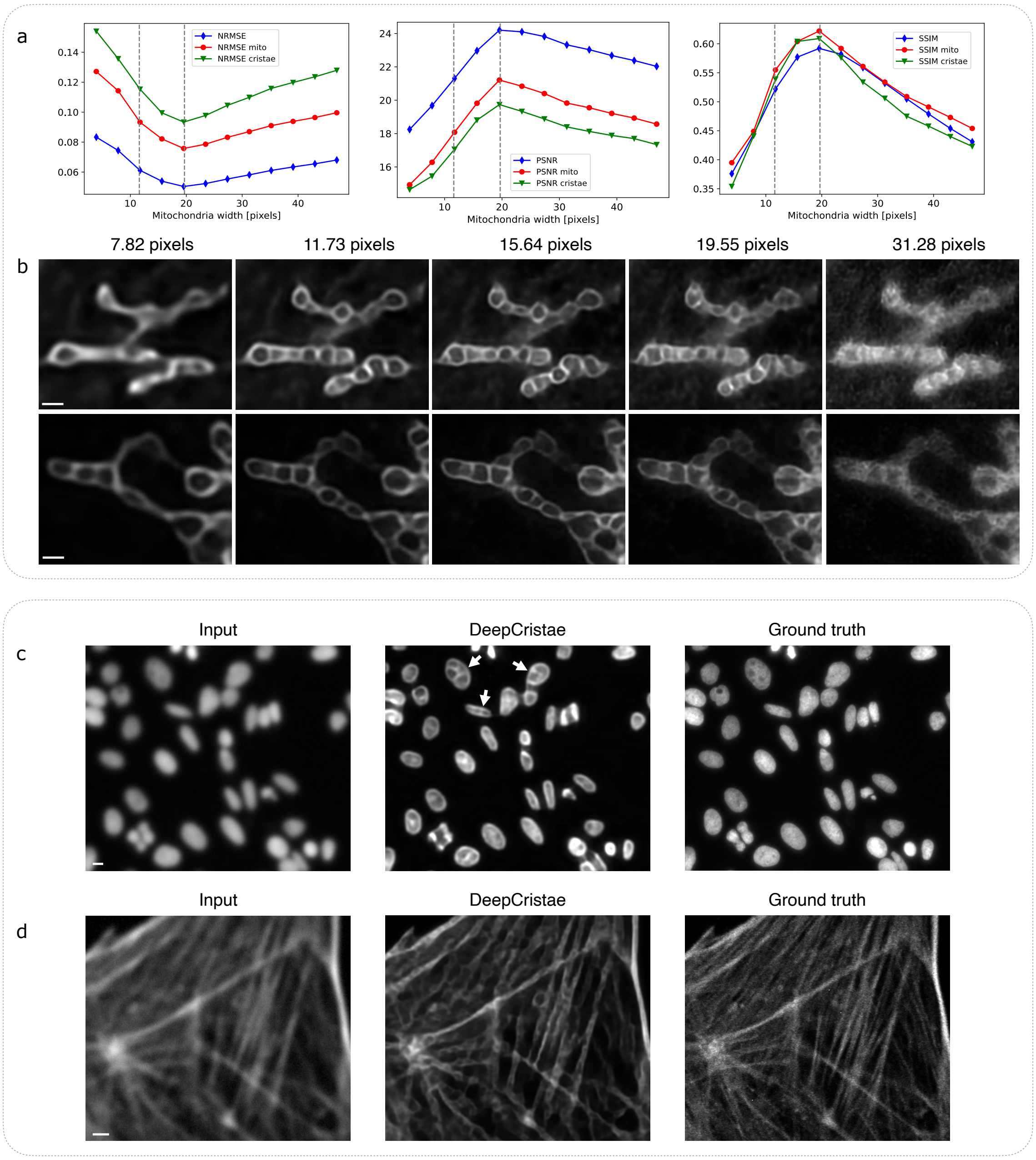


**Supplementary Figure 2. Robustness of DeepCristae to mitochondria scale in LR images and limitations of use.** *First experiment* ***(a, b)***: assessment of the robustness of DeepCristae to the mitochondria width in the image. To that end, DeepCristae was successively applied on different rescaled versions of the test set of $D_{synt}$. Our model was trained from the training set of $D_{synt}$ containing mitochondria of width $15.64 \pm 4.04$ pixels on average**.** **(a)** Evolution of the metrics according to the average width of the mitochondria (in pixels) contained in the test data**.** The two dashed lines on the plots depict the range of mitochondria width contained in the dataset used to train our model. **(b)** Predictions on two test images for five different scaling (left to right: mitochondria of width 7.82, 11.73, 15.64, 19.55, 31.28 pixels on average). Pixel size: 25 nm. Scale bar: 0.5 µm. *Second experiment* **(c-d)**: the use of DeepCristae on applications it was not trained for leads to invalid results. The restoration of images of nucleus[15] **(c)** and actin filaments[16] **(d)** by DeepCristae presents erroneous structures that resemble cristae (white arrows in **(c)**). **(c)** Pixel size: 645 nm. Scale bar: 10 µm**.** **(d)** Pixel size: 96 nm. Scale bar: 2 µm.

a

| | NRMSE | $NRMSE_{mito}$ | $NRMSE_{cristae}$ | PSNR | $PSNR_{mito}$ | $PSNR_{cristae}$ | SSIM | $SSIM_{mito}$ | $SSIM_{cristae}$ |
|---|---|---|---|---|---|---|---|---|---|
| Mean | $0.060 \pm 0.001$ | $0.090 \pm 0.001$ | $0.115 \pm 0.002$ | $22.58 \pm 0.20$ | $19.38 \pm 0.18$ | $17.77 \pm 0.18$ | $0.48 \pm 0.01$ | $0.52 \pm 0.01$ | $0.50 \pm 0.01$ |

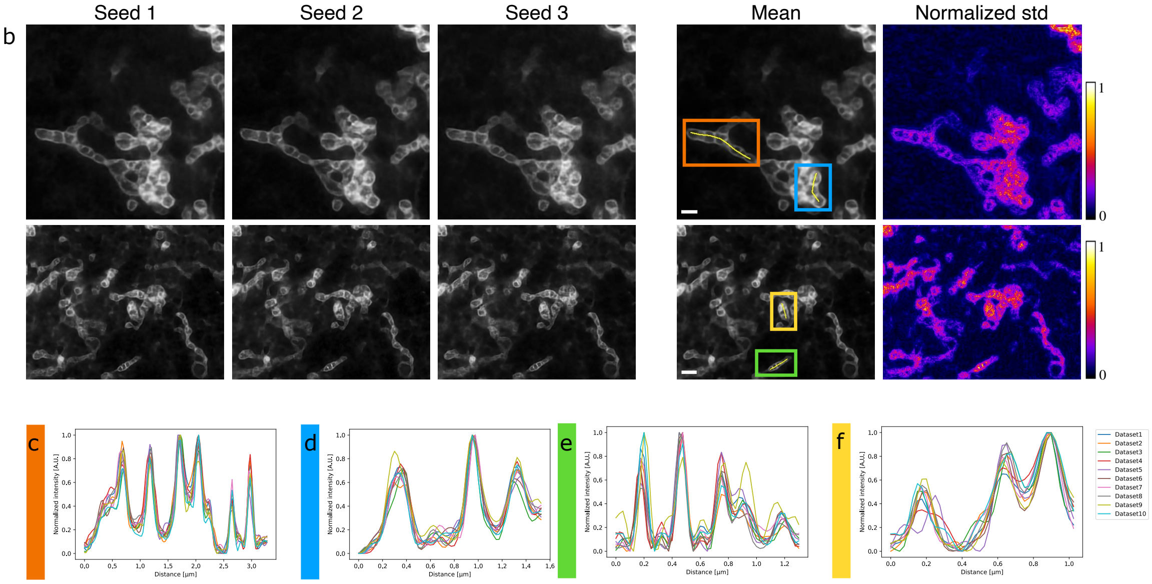


**Supplementary Figure 3. Stability of image restoration by DeepCristae.** Assessment of the reliability of DeepCristae by studying the consistency between its predictions obtained with different trainings. To that end, 10 DeepCristae neural networks were trained with the same training data but with 10 different initializations of weights. This method allows us to conclude whether the same dataset leads to homogeneous learning, meaning that the randomness of initialization does not play a role in the learning process**.** **(a)** Quantitative comparison of the 10 DeepCristae models. Metrics were computed on the test dataset of $D_{synt}$. **(b)** From left to right: predictions of three DeepCristae networks on two images, the average prediction over the 10 trainings and the corresponding pixel-wise normalized standard deviation. Pixel size: 25 nm. Scale bar: 1 µm. **(c-f)** Comparison of normalized intensity line profiles along a mitochondrion between the 10 trainings. The yellow line indicated in the corresponding colored inset in **(b)** serves to identify the fluorescence profile.

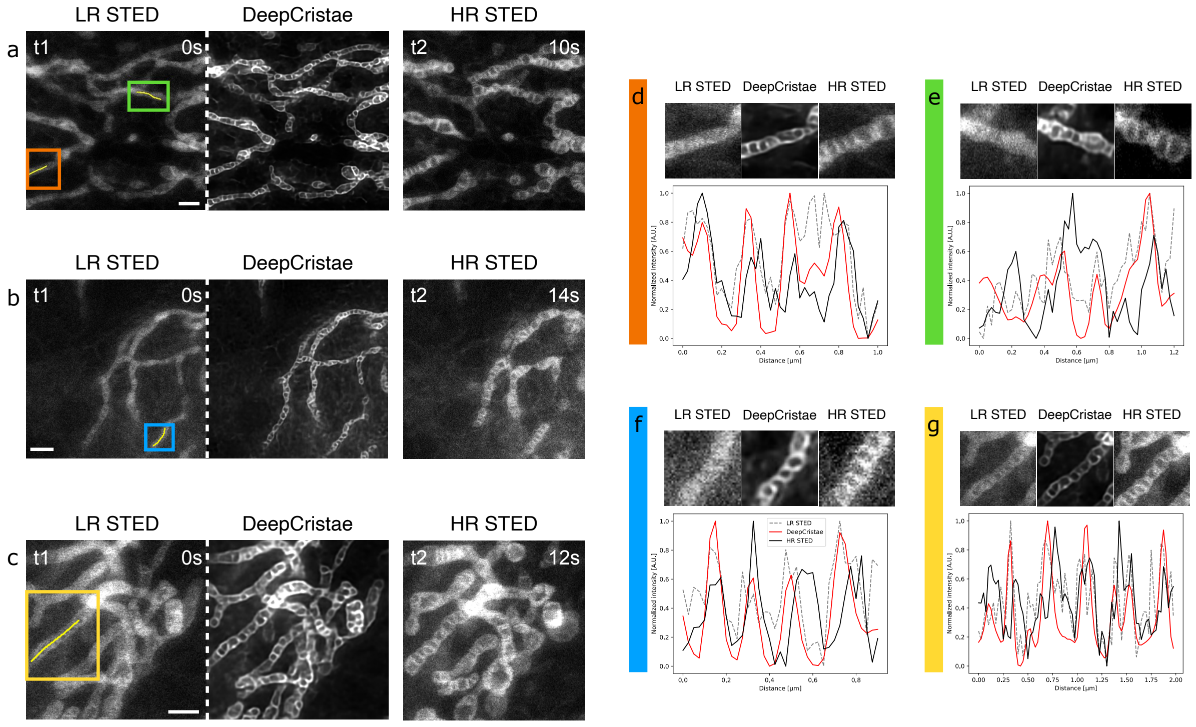


h

| | Mean [nm] | SD [nm] |
|---|---|---|
| DeepCristae | 250,76 | 87,58 |
| HR STED | 243,86 | 84,21 |
| | t-test p-val | F-test p-val |
| DeepCristae - HR STED | ns | ns |

**Supplementary Figure 4. Reliability of image restoration by DeepCristae.** Assessment of the reliability of DeepCristae on real data by controlling the consistency between DeepCristae restoration and a real reference image. RPE1 cells were labeled with PKMITO-Orange prior to low-resolution (LR) and to high-resolution (HR) 2D STED imaging, acquired successively (~30 s of delay). Due to the delay, there are displacements and deformations of mitochondria between the two acquisitions. **(a-c)** Three LR STED images were given as input to DeepCristae for restoration and qualitatively compared to the corresponding HR STED images. Pixel size: 25 nm. Scale bar: 1 μm. **(d)**, **(e)**, **(f)** and **(g)** Zoom in on the colored insets depicted in **(a)**, **(b)** and **(c)**, respectively. Top, from left to right: thumbnails of the zoomed-in area on the LR image, the image restored by DeepCristae and the HR STED image, respectively. Bottom: comparison of normalized intensity line profiles along a mitochondrion between the three thumbnails. The yellow line indicated in the corresponding colored inset in **(a)**, **(b)** and **(c)** serves to identify the fluorescence profile. Note that the line profiles were taken in regions where small mitochondrial displacements were observed. Despite an offset caused by mitochondrial motion, the cristae restored by DeepCristae show overall consistency with those observed in the HR STED images. **(h)** Mean and standard deviation (SD) of the crista-to-crista distance, measured as peak-to-peak intervals (33 intervals for DeepCristae and 35 intervals for HR STED from ten-line plots over six images). The statistical significance Student- and Fisher-tests are also provided; ns= non-significant.

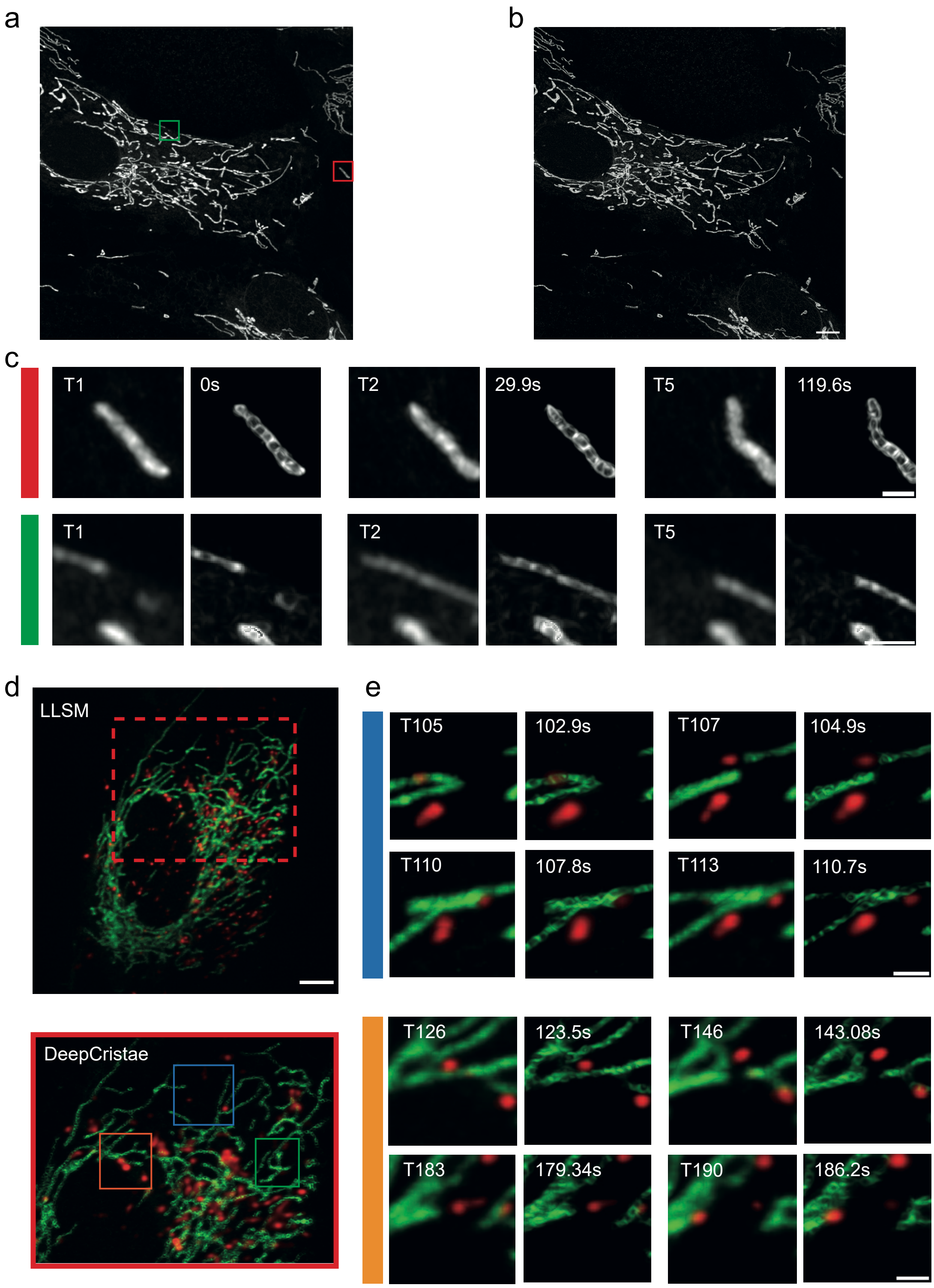


**Supplementary Figure 5. DeepCristae applied to AiryScan processed images and Lattice Light Sheet Microscopy. (a-c)** RPE1 cells were incubated with PKMITO-Orange (gray) for 15 minutes. A MIP image of a 3D stack (15 planes; stack time = 29.9 s/channel, time point T1 out of 5) acquired with an AiryScan microscope is shown before **(a)** and after **(b)** DeepCristae restoration. **(c)** Thumbnails show selected time points of the zoomed area, as indicated by the insets in the red and green squares in **(a)**. They are presented as paired images before (left panels) and after (right panels) DeepCristae restoration. Both time points (left panels) and time frames in seconds (right panels) are indicated for comparison. **(d-e)** RPE1 cells incubated for 4 hours with Cell Mask Plasma Membrane Deep Red (red) were labeled with PKMITO-Orange (green) for the last 15 minutes. **(d)** A maximum intensity projection (MIP) (56 planes; stack time = 0.49 s/channel, time point T1 out of 200) of data acquired with Lattice Light Sheet Microscopy (LLSM) after deskewing and Richardson-Lucy deconvolution is shown before (top) and after (bottom) DeepCristae restoration (green channel). Colored insets indicate intracellular locations with a focus on dynamic events of interest. The upper image shows the Lattice Light Sheet MIP image, while the red square represents the region of interest (ROI) in the DeepCristae image below. **(e)** Thumbnails show selected time points of the zoomed area, as indicated by insets in **(d-bottom)**. Time points are shown as paired images, with the left one before DeepCristae restoration (green channel) and the right one after. Note that for the “blue” area, an early time series (T105-T113) is shown, depicting mitochondria fusion. Similarly, the “orange” area (T126-T190) shows mitochondria fission. Full acquisition video (T1-T200), including these areas as well as the “green” area, which shows a static endosome, is provided as Movie S4. Note that a rescaling factor of 4.16 was applied to the raw LLSM data, and a rescaling factor of 1.68 was applied to the raw AiryScan data before DeepCristae restoration. This is to comply with the conditions for using DeepCristae (see Results, “Robustness of DeepCristae with respect to noise, blur and mitochondria scale in low-resolution images”). Scale bars in **(b, d)** and **(c, e)** are 5 µm and 1 µm, respectively.

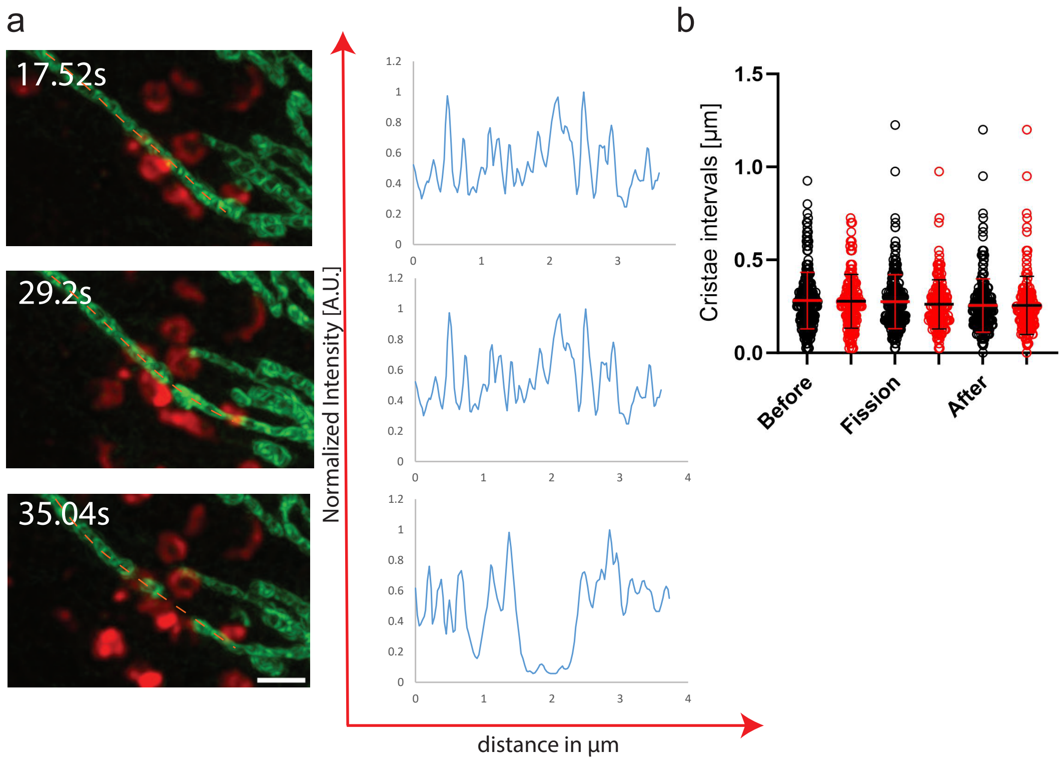


c

| | Cristae intervals | | | Cristae intervals when lysosome contacts | | |
|---|---|---|---|---|---|---|
| | mean [µm] | SD [µm] | N | mean [µm] | SD [µm] | N |
| Before | 0.280 | 0.150 | 422 | 0.277 | 0.144 | 173 |
| Fission | 0.275 | 0.144 | 275 | 0.261 | 0.131 | 161 |
| After | 0.255 | 0.142 | 276 | 0.252 | 0.155 | 161 |

| | t-test p-val | F-test p-val |
|---|---|---|
| Before- Fission | 0.2457[ns] | 0.0006[***] |
| Before- After | 0.0019[**] | <0.0001[****] |

**Supplementary Figure 6. Cristae density evolution during fission in or out of lysosome contacts with mitochondria.** RPE1 cells were incubated with SIR-Lysosome (red) for 2 hours, followed by labeling with PKMITO-Orange (green) for 15 minutes. **(a-left)** Thumbnails show selected time points from a zoomed-in area, displayed as Maximum Intensity Projections (6 planes; stack time = 2.92 s/channel) from images acquired with Live-SR microscopy after DeepCristae restoration (green channel) and following denoising (ND-SAFIR) and Richardson-Lucy deconvolution for lysosome labeling (red channel). The images represent a location where mitochondria fission is occurring. **(a-right)** PKMITO-Orange intensity line plots after DeepCristae. Profile lines are indicated in orange **(a-left)**. **(b)** Graphs display the "peak-to-peak" intervals between cristae in DeepCristae restored images, measured before, during, and after fission. Measurements were first taken from 32 distinct time series, in a blinded manner (black circles; the numbers of peak-to-peak intervals (N) are indicated in **c**), and then from the same series, where lysosome contacts with mitochondria occur (19 of 32 distinct time series; red circles). Error bars indicate mean ± standard deviation (SD). **(c)** Statistics table for the blinded measurements, including mean, SD, and significance levels using Student's and Fisher's tests. Scale bar = 1 µm. Note: a rescaling factor of 2.6 was applied to the raw Live-SR data before DeepCristae restoration. This is to respect the condition of use of DeepCristae (see Results, "Robustness of DeepCristae with respect to noise, blur and mitochondria scale in the low-resolution images").

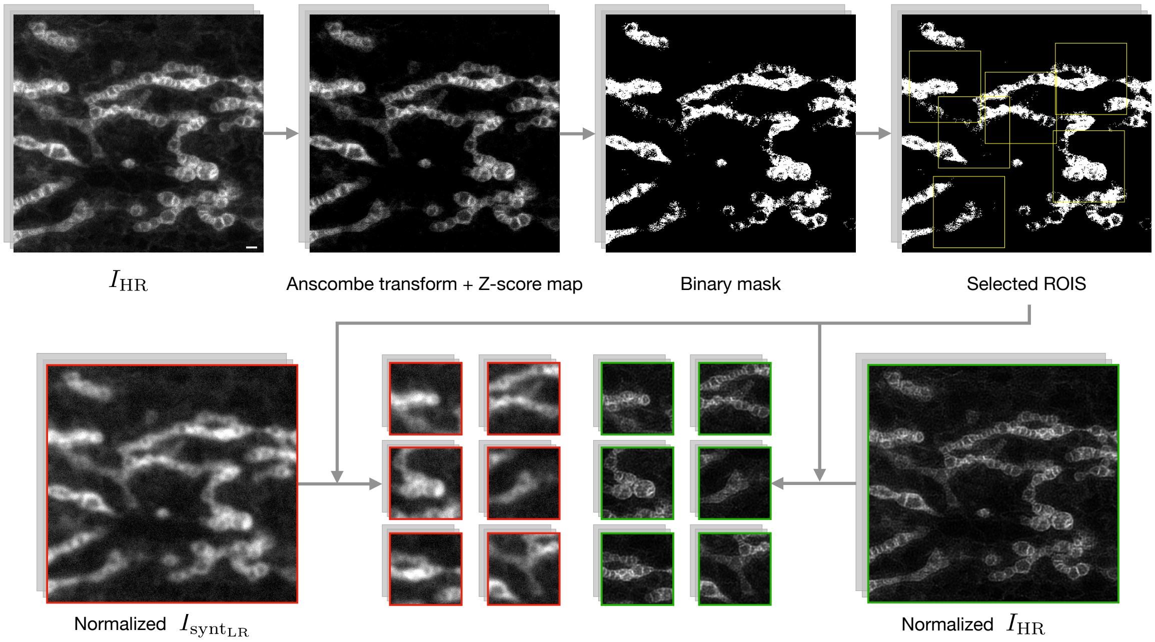


**Supplementary Figure 7. Pipeline of the image patch sampling for the training step.** First, for each HR STED image $I_{HR}$ of the training set of $D_{synt}$, the mitochondria signal is enhanced with respect to the noise. To do this, an Anscombe transform followed by a Z-score map (see Methods, Eq. (4)) is computed. The higher the Z-score, the farther the pixel value is above the mean of the measured noise and therefore considered as a signal of interest. Then, a threshold is applied to the Z-score map to obtain a binary mask of the mitochondria. The value of the threshold is set automatically. Starting from a fixed high value of 30, while the selected pixels do not contain a minimum of 10% mitochondrial information, we subtract 5 from the threshold value. From the obtained binary mask, $N_I$ pixels (in the illustration $N_I$= 6) are chosen to be the center of $N_I$ ROIs of size 128×128 pixels. A minimum distance of 60 pixels is established between each pairwise ROI center. The resulting ROIs are finally used to create the patches from the normalized training data ($I_{synt_{LR}}$ (red) and $I_{HR}$(green) normalized using a percentile normalization). Pixel size: 25 nm. Scale bar: 0.5 µm.

a

| | NRMSE | $\text{NRMSE}_{\text{mito}}$ | $\text{NRMSE}_{\text{cristae}}$ | PSNR | $\text{PSNR}_{\text{mito}}$ | $\text{PSNR}_{\text{cristae}}$ | SSIM | $\text{SSIM}_{\text{mito}}$ | $\text{SSIM}_{\text{cristae}}$ |
|---|---|---|---|---|---|---|---|---|---|
| DeepCristae | 0.059 ± 0.016 | 0.090 ± 0.021 | **0.113 ± 0.026** | 22.75 ± 2.81 | **19.46 ± 2.50** | 17.91 ± 2.30 | **0.50 ± 0.11** | **0.52 ± 0.10** | **0.51 ± 0.15** |
| Depth 2 | 0.063 ± 0.018 | 0.097 ± 0.023 | 0.123 ± 0.027 | 22.31 ± 2.74 | 18.92 ± 2.14 | 17.33 ± 2.03 | 0.47 ± 0.11 | 0.48 ± 0.10 | 0.47 ± 0.15 |
| $\text{SCoP}_{1,1}$ =DSSIM | 0.059 ± 0.017 | 0.089 ± 0.020 | **0.113 ± 0.026** | 22.48 ± 2.58 | 19.44 ± 2.29 | **18.00 ± 2.13** | 0.47 ± 0.10 | **0.52 ± 0.10** | **0.51 ± 0.14** |
| $\text{SCoP}_{1,6}$ | 0.061 ± 0.017 | 0.093 ± 0.021 | 0.117 ± 0.026 | 22.81 ± 2.74 | 19.39 ± 2.11 | 17.72 ± 2.01 | 0.49 ± 0.12 | 0.51 ± 0.09 | 0.50 ± 0.14 |
| MAE | **0.058 ± 0.017** | **0.088 ± 0.021** | 0.114 ± 0.027 | **22.85 ± 2.79** | **19.46 ± 2.75** | 17.86 ± 2.39 | **0.50 ± 0.11** | **0.52 ± 0.10** | 0.49 ± 0.14 |
| No DA | 0.059 ± 0.016 | **0.088 ± 0.019** | **0.113 ± 0.025** | 22.34 ± 2.36 | 19.29 ± 2.16 | 17.91 ± 2.08 | 0.47 ± 0.10 | **0.52 ± 0.08** | **0.51 ± 0.13** |
| Grid | 0.060 ± 0.016 | 0.090 ± 0.019 | **0.113 ± 0.023** | 22.37 ± 2.67 | 19.11 ± 2.11 | 17.75 ± 2.06 | 0.49 ± 0.11 | **0.52 ± 0.09** | **0.51 ± 0.13** |

b

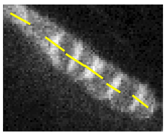

c

| | Cristae width [nm] | Nb cristae detected [%] |
|---|---|---|
| DeepCristae | **137,62 ± 59,64** | **89,03** |
| $\text{SCoP}_{1,1}$ | 148,55 ± 75,56 | 87,10 |
| MAE | 154,31 ± 90,39 | 86,45 |
| No DA | 153,30 ± 66,51 | 87,74 |
| Grid | 153,60 ± 72,49 | **89,03** |
| HR STED | 92,44 ± 23,59 | |

d

| | t-test p-val | F-test p-val |
|---|---|---|
| DeepCristae - $\text{SCoP}_{1,1}$ | ns | 0,0021 [**] |
| DeepCristae - MAE | ns | <0,0001 [****] |
| DeepCristae - No DA | 0,041 [*] | ns |
| DeepCristae - Grid | 0,046 [*] | 0,0115 [*] |

**Supplementary Figure 8. Ablation study for different components of our method.** Only one component is suppressed or modified at a time, respectively: reduction of the size of our network (Depth 2), substitution of our *SCoP* loss by another loss ($SCoP_{1,1}$, $SCoP_{1,6}$, *MAE*), removal of the data augmentation (No DA) and substitution of our patch sampling method by a simple grid sampling one (Grid). Note that the $SCoP_{1,1}$ loss is equivalent to the structural dissimilarity (*DSSIM*) loss **(a)** Quantitative comparison of DeepCristae with its variations. Metrics were computed on the test set of $D_{synt}$. **(b-d)** Measurement of cristae widths for 155 cristae from the 26 test images after restoration obtained with DeepCristae, MAE, $SCoP_{1,1}$, No DA and Grid. Line profiles (as depicted in **(b)**) were fitted to a Gaussian model and FWHM was measured (Supplementary Note 1.3). **(c)** Table with the number of cristae restored by each method, in comparison to the 155 observed in HR STED images, and their average width. **(d)** Table with the statistical significance Student- and Fisher-tests; ns= non-significant.

### Movies

**Movie S1. DeepCristae restoration enhances cristae resolution in live imaging using a Spinning-Disk confocal microscope equipped with Live-SR (related to Fig. 7e).** 3D stacks of live RPE1 cells labeled with PKMITO-Orange (also known as PKMO) were acquired with a Spinning-Disk confocal microscope equipped with a Super Resolution module. Mitochondrial fission events are shown.

**Movie S2. DeepCristae restoration enhances cristae resolution in live imaging using a Lattice Light Sheet microscope (related to Fig. 7f).** 3D stacks of live RPE1 cells labeled with PKMITO-Orange (also known as PKMO) were acquired with a Lattice Light Sheet Microscope (denoted LLSM). Fusion and fission dynamics of mitochondria are indicated in a zoomed region.

**Movie S3. DeepCristae reveals 3D+time cristae morphology during endo/lysosome mitochondria interactions using a Spinning-Disk confocal microscope equipped with Live-SR (related to Fig. 8a).** RPE1 cells incubated with CellMask Plasma Membrane Deep Red (red) (denoted PMDR) were labeled with PKMITO-Orange (also known as PKMO) (green). 3D stacks were acquired every 1.86 s per channel with Live-SR microscopy. Data is shown before and after DeepCristae restoration of the mitochondria (green channel) as well as after denoising (ND-SAFIR) and Richardson-Lucy deconvolution of the endo/lysosomes (red channel). Insets correspond to Fig. 8a.

**Movie S4. DeepCristae reveals 3D+time cristae morphology during endo/lysosome mitochondria interactions using Lattice Light Sheet microscope (related to Supplementary Fig. 5d-e).** RPE1 cells incubated with CellMask Plasma Membrane Deep Red (denoted PMDR) (red) were labeled with PKMITO-Orange (also known as PKMO) (green). 3D stacks were acquired every 0.49 s per channel with Lattice Light Sheet Microscopy (denoted LLSM). Data is shown after deskewing and Richardson-Lucy deconvolution before and after DeepCristae restoration of the mitochondria (green channel) as well as after denoising (ND-SAFIR) of the endo/lysosomes (red channel). Insets correspond to Supplementary Fig. 5d.
